# Co-delivery of neurotrophic factors and a zinc chelator substantially promotes axon regeneration in the optic nerve crush model

**DOI:** 10.1101/2024.11.20.624564

**Authors:** Huynh Quang Dieu Nguyen, Mi-hyun Nam, Jozsef Vigh, Joseph Brzezinski, Lucas Duncan, Daewon Park

## Abstract

Traumatic optic neuropathies cause the death of retinal ganglion cells (RGCs) and axon degeneration. This is a result of the blockage of neurotrophic factor (NTF) supply from the brain and a vicious cycle of neurotoxicity, possibly mediated by increased levels of retinal Zn^2+^. Ciliary neurotrophic factor (CNTF) and brain-derived neurotrophic factor (BDNF) are two NTFs that are known to support RGC survival and promote axon regeneration. Dipicolylamine (DPA) has a strong affinity to Zn^2+^ and can selectively chelate this ion. To continuously supply NTFs and reduce elevated retinal Zn^2+^, we developed poly(serinol hexamethylene urea)-based sulfonated nanoparticles (S-PSHU NPs), that co-delivers CNTF, BDNF, and DPA. An *in vitro* release study was performed using the NTF-DPA-loaded S-PSHU NPs, demonstrating a sustained release of CNTF and BDNF for up to 8 weeks, while DPA was released for 4 weeks. In a rat optic nerve crush (ONC) model, DPA-loaded S-PSHU NPs exhibited dose-dependent elimination of retinal Zn^2+^. Similarly, *in vitro* primary RGC culture demonstrated that the activity of RGCs and axon growth were dependent on the dosage of CNTF and BDNF. In addition, the NTF-DPA-loaded S-PSHU NPs significantly improved RGC survival and axon regeneration following ONC in rats, with the regenerated axons extending to the distal segment of the brain, including the suprachiasmatic nucleus, lateral geniculate nucleus, and superior colliculus.

## INTRODUCTION

Traumatic optic neuropathy causes the death of retinal ganglion cells (RGCs) leading to vision loss [1]. This optic neuropathy causes the obstruction of axonal transport of neurotrophic factors (NTFs). Consequently, many treatment approaches have focused on delivering NTFs directly into the vitreous humor in various animal models [2–4]. Previously, we reported that the delivery of NTFs significantly increases RGC survival and axon extension in a rat model [5].

While the effects of NTFs in RGCs protection and axon regeneration are evident, the therapeutic use of NTFs has been greatly limited. NTFs do not cross the blood–brain barrier and their effects are transient due to short *in vivo* half-lives by rapid inactivation [6]. To continuously supply NTFs, various strategies include: injecting native NTFs [3, 6], transplanting NTF- secreting cells [7, 8] and peripheral nerve (PN) grafts [9, 10]. These methods fail to achieve clinical success due to their intrinsic drawbacks, including trauma risks due to frequent injections, low survival rate of transplanted cells, and limited NTF delivery from PN grafts.

Traumatic optic neuropathy also results in neurodegeneration caused by a vicious cycle of neurotoxicity [11], due to which solely supplying NTFs may limit RGC survival and axon regeneration. For decades, researchers have believed that pro-inflammatory reactions drive neurotoxicity [11–13]. However, inhibition of the pro-inflammatory reaction has not resulted in success in clinical applications. Recently, a study reported that the intraocular chelation of mobile zinc (Zn^2+^) using a *N,N,N′,N′*-tetrakis (2-pyridylmethyl) ethylenediamine (TPEN) greatly improved RGC survival and axon regeneration after the optic nerve injury [14]. However, even after the early administration of TPEN, the continued low-level accumulation of Zn^2+^ sufficient to cause RGC death was observed. Additional TPEN administration 4 and 7 days after the first dose was also more effective than single administration.

Based on these facts, a system that can effectively eliminate the elevated retinal Zn^2+^ while continuously supporting NTFs is a good combinatorial strategy to increase RGC survival and axon regeneration after traumatic optic neuropathy. To achieve such a system, we created nanoparticles (NPs) loaded with NTFs and a Zn^2+^ chelator. The NPs were fabricated using a sulfonated poly(serinol hexamethylene urea) (S-PSHU) [15].

Ciliary neurotrophic factor (CNTF) and brain-derived neurotrophic factor (BDNF) were utilized as NTFs. BDNF is a powerful neurotrophin for developing and adult RGCs [16]. Studies reported that BDNF administration enhances RGC survival in optic nerve crush models [17, 18]. Thus, BDNF may play a crucial role in protecting RGCs from optic nerve degeneration; however, it is also known to have limited effect on RGC axon regeneration [19]. In the retina, mRNA expression of CNTF increases after optic nerve injuries; in concert, RGCs express high affinity CNTFRα receptors [20, 21], suggesting that CNTF is an important factor for RGC survival. Indeed, CNTF-based treatment enhanced RGC survival in ONC models [3, 22]. Most importantly, unlike BDNF, CNTF stimulates RGC axon regeneration after optic nerve injuries [16], as we observed previously [5]. Thus, the co-delivery of BDNF and CNTF may increase RGC survival and promote RGC axon regeneration.

Dipicolylamine (DPA) was employed as the Zn^2+^ chelator. DPA selectively chelates zinc ions [23] via a strong affinity to zinc (Kd ∼ 10^-11^ M) [24–26]. DPA does not associate strongly with most metal ions, which exist at high concentrations under a physiological condition [24, 27]. The selectivity of DPA to zinc has been well documented: DPA-based fluorescent probes (e.g., Newport Green) selectively labels labile zinc in viable beta-cells [28], is used to measure zinc binding to amyloid-beta (Aβ) [29] and has been used to quantitatively assess zinc concentration under various physiological conditions [30–34].

After loading CNTF, BDNF, and DPA in the S-PSHU NPs, the efficiency of this delivery system was evaluated by establishing the release profiles and determining RGC growth *in vitro*. Further, the treatment efficiency was also evaluated by an intravitreal injection of the system in a rat optic nerve crush model. Overall, the co-delivery of CNTF, BDNF, and DPA showed great potential for the treatment of traumatic optic neuropathy with enhanced RGC protection and extensive axon regeneration into the visual target of the brain.

## MATERIALS AND METHODS

### Materials

CNTF, BDNF, MTT assay kit, CNTF and BDNF ELISA kits were purchased from Peprotech (Rocky Hill, NJ, USA). N-BOC-serinol, hexamethylene isocyanate, urea, bovine serum albumin (BSA), Zinpyr-1, DPA, dimethyl sulfoxide (DMSO), acetonitrile, and phosphoric acid were purchased from Sigma Aldrich (St. Louis, MO). Xylazine and ketamine were purchased from MWI Veterinary Supply (Boise, ID). Wistar rats (female, 200-250 g) were purchased from Charles River Laboratories (Wilmington, MA). Beta III tubulin (Tuj-1, MA1-118), Alexa Fluor 555 conjugated cholera toxin subunit B (CTB, C34776), GAP-43 (33-5000), and Phosphate Buffered Saline (PBS) were purchased from Life Technologies (Carlsbad, CA).

### Equipment

Tissue was sectioned using a CryoStar NX70 Cryostat. Confocal images were obtained using a Nikon Eclipse Ti C2 LUN-A microscope (Nikon, Tokyo) equipped with two C2-DU3 high sensitivity PMT detectors, and a motorized microscope stage with 3-axis navigation (X, Y, and Z). Fourier transform infrared spectroscopy (FTIR) was performed on a Nicolet 6700 FTIR Spectrometer using polyethylene infrared sample cards. Scanning electron microscope (SEM) images were taken using a JEOL (Peabody, MA) JSAM-6010la. ELISA color development was measured by a plate reader (BioTek Synergy 2 Multi-Mode Reader) at 405 nm with wavelength correction set at 650 nm. DPA was quantitated by HPLC (Agilent 1220) equipped with a Venusil C18 column (VA952505-0), an autosampler and a visible light absorbance detector.

### Fabricate S-PSHU NPs loaded with three agents and determine release profiles

As the material used to formulate the NPs, S-PSHU was synthesized and characterized as we described previously [15, 35]. Briefly, PSHU was synthesized using N-BOC-serinol, hexamethylene diisocyanate, and urea. Subsequently, the amine groups (–NH2) in PSHU were sulfonated using a propane sultone to synthesize S-PSHU.

For the fabrication of NTF-DPA-loaded S-PSHU NPs, 15 µg of S-PSHU, 1 µg each of CNTF and BDNF and 15 µg DPA were dissolved in 20 µl DMSO. Subsequently, the solution was added dropwise to 20 mL of milliQ water for 10 min while sonicating. The S-PSHU NPs, loaded with each of the three agents, were recovered by centrifugation at 14,000 rpm for 15 min, pouring off the supernatant and then re-suspending the NPs in milliQ water. This procedure was carried out three times. Please note that we followed the same protocol to prepare agents-loaded NPs throughout the studies unless described specifically. For comparison, the NTF-DPA-loaded PSHU-NPs without sulfonate groups were also prepared using the same procedure.

To determine the release profiles, as-prepared NTF-DPA-loaded S-PSHU NPs were dispersed in 1 ml of PBS containing 0.1 % BSA in a 2.5 ml centrifuge tube. Then, the suspension was placed on an orbital shaker with gentle shaking at 37°C. At predetermined time points, 1 ml of supernatant was withdrawn after centrifugation, and then refreshed with another 1 ml of PBS-BSA. For each of the 3 agents, 300 µl of the supernatant sample was used for quantification.

After resuspension, this process was repeated until the end point of the release test was reached. The levels of CNTF and BDNF were quantified using agent-specific ELISA kits. DPA was quantified using HPLC with a reverse phase column and mobile phase of acetonitrile/water/phosphoric acid (20/80/0.1) at a flow rate of 1 ml/min.

### Optic nerve crush (ONC) model

All animal experiments were performed in accordance with procedures approved by Animal Care and Use Committees at the University of Colorado Denver Anschutz Medical Campus and follow the NIH Guide for the Care and Use of Laboratory Animals. In designing the study, all efforts were made to replace, reduce or refine in accordance with the 3R principle.

Wistar rats (female, 200-250 g) were anesthetized by intraperitoneal injection of a solution (1 ml/kg) containing 10 mg/kg xylazine and 80 mg/kg ketamine. Then, a lateral canthotomy was performed after anesthetizing the eye with proparicaine and then sterilizing the area surrounding the lateral corner of the eye with betadine. The eye was then irrigated using normal saline. Using a straight hemostat, the skin at the lateral corner of the eye was crimped all the way down to the orbit for 1 minute (to achieve hemostasis). The skin around the lateral orbit was lifted with forceps and a 0.5 cm incision was made with scissors. A blunt dissection was utilized to expose the optic nerve. Using a number 5 Jeweler’s forceps, the optic nerve was crimped three times for 10 seconds each approximately 1.5 mm behind the globe [5].

### Preparation of optic nerve and retinal tissue sections

At each pre-determined time points, animals were euthanized using an unchanged cage and a flow rate of CO2 introduced to displace 20% of the cage volume per minute. A bilateral thoracotomy was performed as a secondary method of euthanasia. Eyes were then enucleated and fixed using 4 % PFA in PBS for 30 min. After removing the cornea and lens, the tissues were post-fixed using 4% PFA for 30 min. The tissues were then cryoprotected with 10%, 20%, and 30 % sucrose in PBS for a total of 2 days, embedded in optimal cutting temperature (OCT) compound, and frozen at -80 °C. Longitudinal tissue sections for both analysis of the optic nerve (10 μm thickness) and the retina (5 μm thickness) were collected along the nasal-temporal plane.

### Preparation of whole-mounted retina

For the retina flatmount analysis, four radical cuts were performed from the retinal cup to the equator. Then, the layer of RPE, choroid, and sclera was carefully removed. The remaining retinal cups were washed in 1× PBS thrice.

### Preparation of brain sections

Animals were perfused as described above. Brains were carefully removed and post-fixed in 4% PFA at 4°C overnight. After post-fixation, brains were washed in PBS for 30 minutes, cryoprotected in a graded series of sucrose solutions, embedded in OCT compound, and frozen at -80 °C. The coronal sections (50 μm thickness) were obtained in the area of suprachiasmatic nucleus (SCN), lateral geniculate nucleus (LGN), and superior colliculus (SC).

### Determine the Zn^2+^ eliminating capacity of DPA

To test the zinc-chelating capability of DPA, various formulations of S-PSHU NPs were prepared using 15 µg of S-PSHU and varying amounts of DPA (1, 5, 10 and 15 µg), while maintaining CNTF and BDNF each at 1 µg. The retrieved NPs were resuspended in 5 µl of PBS and injected into the vitreous humor of rats using a 32-gauge needle immediately following ONC. To capture Zn^2+^ accumulation, 5 µl aliquot of Zinpyr-1 (ZP-1, 500 µM) was intravitreally injected [14]. One day after the injection, animals were sacrificed, and the retinal tissue sections were obtained as described above. For comparison, the retinal accumulation of Zn^2+^ in healthy and non-treatment groups were also monitored. The intensity of ZP-1 expression was monitored by a confocal microscopy and subsequently quantified with ImageJ software. Composite images of whole cross-sections of the retina were obtained. Z-stack maximum intensity images of 5 total stacks were taken to encompass the full section height (5 μm). The expression of ZP-1 was quantified by determining the average level of fluorescence intensity.

### Determine the loading amounts of CNTF and BDNF

To determine the loading amounts of NTFs, primary RGCs were cultured with different formulations. Primary RGCs were isolated from juvenile Wistar rats (postnatal 5-7) as we described previously (Supplementary material) [35]. Different formulations were prepared by loading varying amounts of CNTF and BDNF (0.2 - 1 µg with 0.2 µg increments). The retrieved NPs were added to 24-well plates containing RGCs (10^4^ cells/well). For the control group, RGCs were cultured in NTF-free culture medium containing MACS NeuroMedium (130-093-570, Miltenyl Biotec, San Diego, CA), NeuroBrew-21 (1:50, 130-093-566, Miltenyl Biotec, San Diego, CA), sodium pyruvate (1 mM), N-acetylcystein (50 μg/ml), insulin (5 μg/ml), Forskolin (10 μM), glutamine (2 mM), triiodothyronin (40 ng/ml), streptomycin sulfate (100 μg/ml), and penicillin (100 U/ml). At predetermined time points, cell proliferation and axon growth were examined by MTT assay and Tuj-1 (1:100) staining, respectively [35]. The average axon length and density were quantified by analyzing the maximal intensity of images, as previously described (Supplementary material) [35].

### Evaluate the bioactivity of the encapsulated NTFs

NTF-loaded S-PSHU NPs and NTF-loaded PSHU NPs were prepared using 15 µg of each polymer, and 0.6 µg each of CNTF and BDNF. The retrieved NPs were dispersed in 1 ml of PBS containing 0.1 % BSA in a 2.5 ml centrifuge tube. Subsequently, the suspension was placed on an orbital shaker with gentle shaking for 4 weeks at 37°C. The suspension was centrifuged to remove all supernatant and refreshed with 1 mL of PBS each week. At 4 weeks, the NPs were recovered by centrifugation and resuspended in 200 μl of NTF-free RGC culture medium [36].

Subsequently, the NP suspension was added to 24-well plates containing RGCs (10^4^ cells/well). For control groups, RGCs were cultured in 1) NTF-supported culture medium (same as NTF-free culture medium but with 25 ng/ml of BDNF and 10 ng/ml of CNTF) [35], 2) NTF-free culture medium containing freshly prepared NTF-loaded S-PSHU NPs, and 3) NTF-free culture medium containing freshly prepared NTF-loaded PSHU NPs. After 7 days of culture, the axonal growth of RGCs were monitored by Tuj-1 (1:100) staining [35], and the length of the axons was quantified using ImageJ software.

### Evaluate RGC survival and axon regeneration *in vivo*

To evaluate the efficacy of NTF-DPA on RGC survival and axonal regeneration, S- PSHU NPs were formulated using 15 µg of S-PSHU, 0.6 µg each of CNTF and BDNF, along with 10 µg of DPA. After ONC, the retrieved NPs were suspended in 5 µl of saline and intravitreally injected. To assess their individual effectiveness, NTF-loaded S-PSHU NPs and DPA-loaded S-PSHU NPs were also employed. As a control, a 5 µl aliquot of saline was intravitreally injected.

After 8 weeks, animals were sacrificed, and the optic nerve sections and the retina flatmounts were prepared as described above. The loss of RGCs was quantified by immunolabeling the retinal flatmount with Tuj-1 (1:100) staining [37, 38]. The number of Tuj-1- stained RGCs per image was determined. Then, the values were averaged per retina. The axonal regeneration was assessed by tracing Alexa Fluor 555-conjugated CTB in the optic nerves [39, 40]. A 2 µl aliquot (1 µg/ml) of CTB was intravitreally injected 72 h prior to sacrifice. The CTB- positive axons in the optic nerve sections were counted manually at different distances (1, 2, 3, 4, and 4.5 mm) from the crush site in at least five longitudinal sections per group. These values were converted into the number of regenerated axons as described previously [41, 42].

The extension of the regenerated axons to the visual target area in the brain was also examined. Brain sections in the area of SCN, LGN and SC were obtained as described above. Composite images of these areas were obtained. Z-stack maximum intensity images of 10 total stacks were taken to encompass the full section height (50 μm). The expression of CTB-positive axons was quantified by determining the average level of fluorescence intensity.

All images were acquired by a confocal microscopy and further quantified using ImageJ software.

## Statistical analysis

Statistical analyses were performed by ANOVA and unpaired Student’s t-tests, with Bonferroni’s adjustment for multiple comparison adjustments. The significance level was set at p<0.05. Data in figures are presented as mean ± SEM.

## RESULTS AND DISCUSSION

### The S-PSHU NPs showed sustained release of CNTF, BDNF, and DPA

To continuously supply NTFs and eliminate retinal Zn^2+^ in optic neuropathies, a NTF- DPA-loaded S-PSHU NPs system was developed. We began by synthesizing PSHU (Mn: 12,743), which was further sulfonated using propane sultone to synthesize S-PSHU (Mn: 17,086) (Figure S1). Based on Mn changes, ∼89 % of –NH2 were conjugated with sulfonate groups. The conjugation of propane sultone was confirmed by observing S=O and S-O stretching at 1,260 cm^-1^ and 799 cm^-1^, respectively, using FTIR (Figure S2). S-PSHU structurally consists of a PSHU backbone and negatively charged sulfonate groups as side chains. The negatively charged sulfonate group mimics the biofunction of natural heparin interactions between the extracellular matrix and NTFs (e.g. heparin interacting with NTFs). This interaction between the sulfonate groups and the positively charged receptor binding site on NTFs can 1) sustain the release of encapsulated NTFs and 2) protect the proteins from proteolytic degradation, preserving their bioactivity [43, 44].

To compare the advantage of the presence of sulfonate groups, NTF-DPA-loaded S- PSHU NPs and NTF-DPA-loaded PSHU NPs were prepared. The recovery process by centrifugation at 14,000 rpm for 15 min consistently resulted in 99.3% yield. The NTF-DPA-loaded S-PSHU NPs showed a relatively uniform diameter of 223±16 nm (Figure S3). We observed a similar diameter of NTF-DPA-loaded PSHU NPs (∼232 nm, images not shown). The loading efficiency was calculated by measuring the amounts of agents present in the initial supernatant (Table 1). The amount of each agent in the second and the third supernatant were negligible.

**Table 1.** Loading efficiency (LE) of CNTF, BDNF, and DPA in S-PSHU NPs and PSHU-NPs.

| | LE (%) ± SD | |
| --- | --- | --- |
|  | S-PSHU NPs | PSFU NPs |
| CNTF | $73 \pm 5$ | $48 \pm 6$ |
| BDNF | $76 \pm 4$ | $43 \pm 3$ |
| DPA | $43 \pm 6$ | $40 \pm 5$ |

S-PSHU NPs could encapsulate CNTF, BDNF, and DPA simultaneously. Both CNTF and BDNF demonstrated similar loading efficiency. However, DPA showed lower loading efficiency than CNTF and BDNF. This might be attributed to the much smaller molecular size of DPA (640 Da) compared to CNTF (22.7 kDa) and BDNF (28 kDa). However, since the total injection volume is 5 µl for *in vivo* study, ∼40% loading efficiency of DPA is equivalent to ∼1.9 mM, which is high concentration for a Zn^2+^ chelator based on prior literature [14]. PSHU NPs could also encapsulate three agents; however, the loading efficiency of CNTF and BDNF was much lower compared to that of the S-PSHU NPs. The absence of sulfonate groups may have contributed to the difference in loading efficiency.

Following fabrication of the NTF-DPA-loaded NPs, the release profiles were determined (Figure 1). Importantly, the release of all agents exhibited sustained patterns. In the S-PSHU NPs group, we observed the initial burst release of all agents, with both CNTF and BDNF exhibiting significantly lower amounts of burst release and more sustained release compared to DPA (Figure 1A). The release of CNTF and BDNF was consistently sustained throughout the entire 8- week observation period. This could be due to the interaction between the negatively charged sulfonate group in S-PSHU NPs and the positively charged domain in CNTF and BDNF. It could also result from the ∼34 – 44 times larger molecular size of the NTFs compared to DPA, delaying diffusion from S-PSHU NPs. These release profiles might be beneficial for the following reasons. First, since the level of retinal Zn^2+^ rapidly increases within 3 days after optic nerve injuries, the initial burst release of DPA may effectively eliminate the acutely accumulated Zn^2+^ at the early stage. DPA released thereafter may deplete the continuously accumulating Zn^2+^. Second, RGC axon regeneration is a long process. For substantial axon regeneration, RGCs need to survive for long periods of time, which may require continuous NTF supply. For comparison, the same agents were loaded in non-sulfonated PSHU NPs, which resulted in faster release of NTFs compared to the S-PSHU NP group, with a comparable release rate of DPA (Figure 1B).

**Figure 1.**
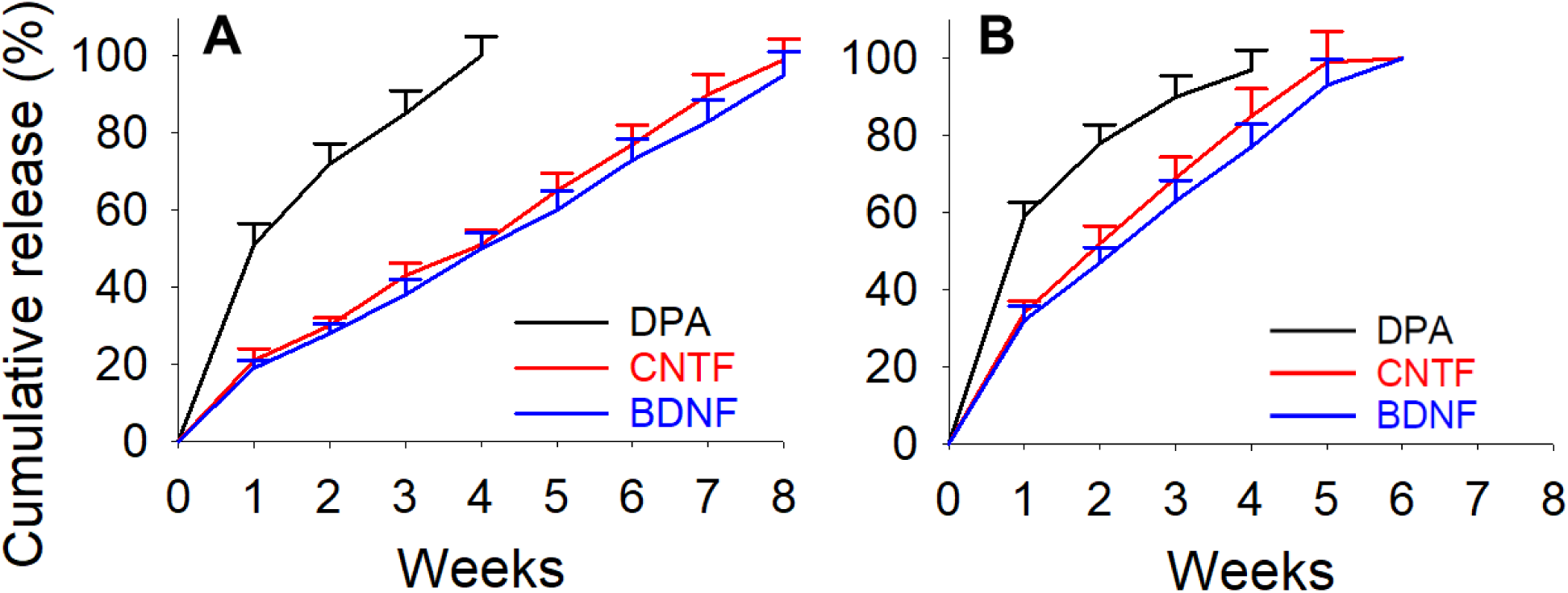
The cumulative release of CNTF, BDNF and DPA from S-SPHU NPs (A) and PSHU NPs (B). All agents showed sustained release profiles. In both groups, DPA was released until 4 weeks, which was much faster than NTFs. The release of NTFs in S-PSHU NPs was sustained until 8 weeks, while the release from PSHU NPs was negligible at 6 weeks. N=5. Error bars indicate standard deviation of the mean.

The absence of sulfonate groups in PSHU NPs might limit the interaction with NTFs and lead to faster release.

### The amounts of retinal Zn^2+^ showed a DPA dose-dependent reduction in the ONC model

To determine the appropriate loading amounts of DPA, various formulations of S-PSHU NPs with varying amounts of DPA (1, 5, 10 and 15 µg) were intravitreally injected in the ONC model. Twenty-four hours after ONC, the zinc accumulation in the no treatment group demonstrated an intensity over 10 times higher than the healthy control (p<0.001), indicating pronounced acute Zn^2+^ accumulation following ONC (Figure 2). This accumulation was particularly notable within the inner nuclear layer (INL) and certain portions of the ganglion cell layer (GCL). In addition, the decrease in ZP-1 intensity exhibited a dose-dependent response to DPA. The intensity significantly decreased up to the 10 µg dosage of DPA-loaded NPs compared to no treatment group (p<0.05, p<0.01, and p<0.001 for 1 µg of DPA, 5 µg of DPA, and 10 µg of DPA, respectively). However, 15 µg of DPA showed no additional decrease in ZP-1 intensity compared to 10 µg of DPA. The ZP-1 intensity of 10 µg- and 15 µg-loaded DPA NPs was similar to that of the healthy control, although it showed slightly increased trends. Therefore, we decided to employ a 10 µg DPA loading into the NPs to further investigate its neuroprotective effects after ONC in rats.

**Figure 2.**
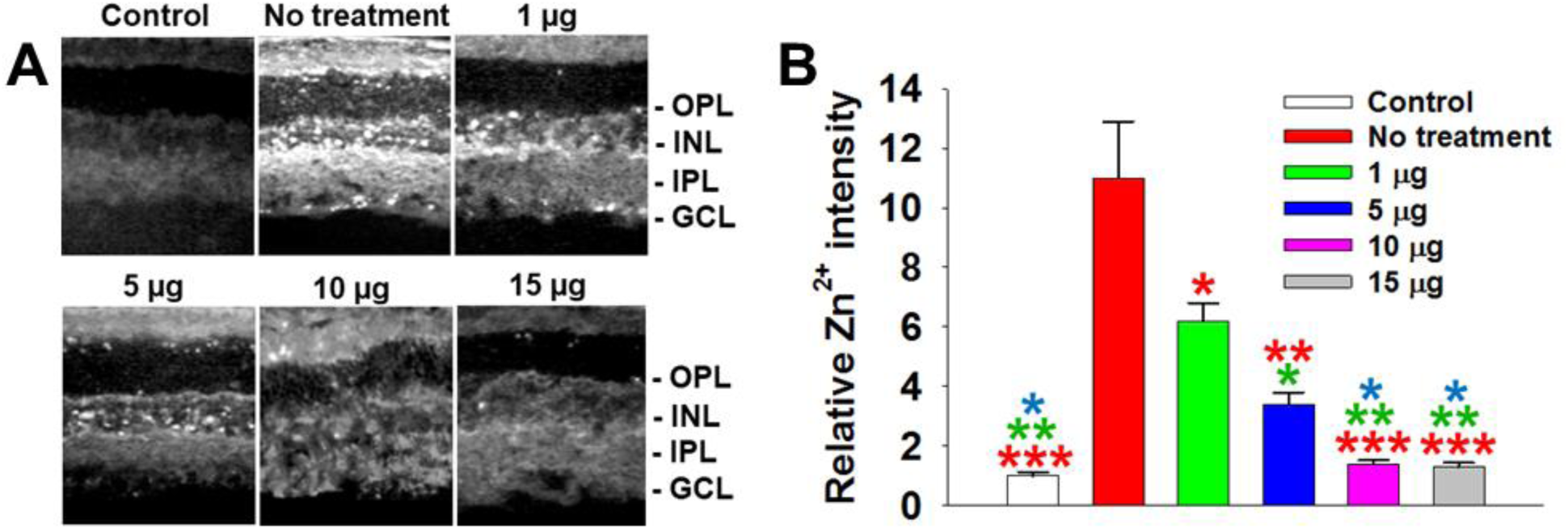
The elimination of Zn^2+^ that accumulated in the retina after ONC. (A) Twenty-four hours after ONC, significant increase in the retinal Zn^2+^ was observed in INL and GCL in no treatment group while treatment with DPA-loaded S-PSHU NPs significantly reduced the level of retinal Zn^2+^. (B) The level of retinal Zn^2+^ was reduced dose-dependently up to 10 µg of DPA; however, no further decrease in Zn^2+^ level was observed thereafter. N=5. *p<0.05, **p<0.01, ***p<0.001 (ANOVA with Bonferroni post-hoc analysis). Error bars indicate standard deviation of the mean.

Zn^2+^ is an essential micronutrient in the nervous system, which plays a crucial role in neural cell proliferation [45–47], neuronal differentiation [48, 49], the modulation of synaptic transmission [50, 51] and cell survival [45, 52]. However, high levels of Zn^2+^ cause clinical problems. Acute neuronal injuries release excessive amounts of Zn^2+^ from terminals or Zn^2+^ binding proteins, and the accumulation of such Zn^2+^ leads to death of cells in the cerebral cortex and hippocampus [56]. In particular, excess Zn^2+^ accumulation in disease conditions has shown toxic effects in retinal ischemia [57, 58], global ischemia [59] and hypoglycemia mediated neuronal death [60, 61]. As evidenced in Figure 2, our NP formulation can eliminate the excess amounts of Zn^2+^ that accumulate in the retinal layer 24 h after ONC. The 24 h time point was chosen as it has been observed that the point of highest level of Zn^2+^ accumulation occurs 24 h post ONC [14]. This is identified as the ideal time point to evaluate the excess Zn^2+^ accumulation and its elimination efficiency by the system. While some Zn^2+^ accumulation was observed in GCL, the most Zn^2+^ accumulated in INL within 24 h post ONC, where horizontal cells, bipolar cells, amacrine cells, and Müller glia reside. The delivery of DPA into the retina effectively eliminated excess Zn^2+^ following ONC. Based on our observation, 10 µg or higher DPA loading seemed to effectively eliminate the excess Zn^2+^ after ONC.

### RGC activity and axon growth were NTF dose-dependent

To determine the loading amount of NTFs, we investigated the effect of various amounts of NTF-loaded NPs on RGC growth. Primary RGCs were cultured with different NTF ratios, each prepared with 15 µg of S-PSHU and varying amounts of NTFs (0.2, 0.4, 0.6, 0.8, and 1.0 µg each of CNTF and BDNF). At 1, 3 and 7 days, RGC activity was measured by MTT assay.

All groups showed a similar trend with increasing activity throughout the observation periods (Figure 3A). In the control group, the activity significantly increased at 3 (p<0.05) and 7 days (p<0.05) of culture. All NTF-loaded S-PSHU NPs resulted in significantly higher activity than the control (p<0.01 for 0.2 µg group and p<0.001 for all other groups). The activity also increased with higher amounts of CNTF and BDNF up to 0.6 µg (p<0.01 for 0.2 µg *vs.* 0.4 µg groups; p<0.05 for 0.4 µg *vs.* 0.6 µg groups). However, no further changes in activity were observed thereafter.

**Figure 3.**
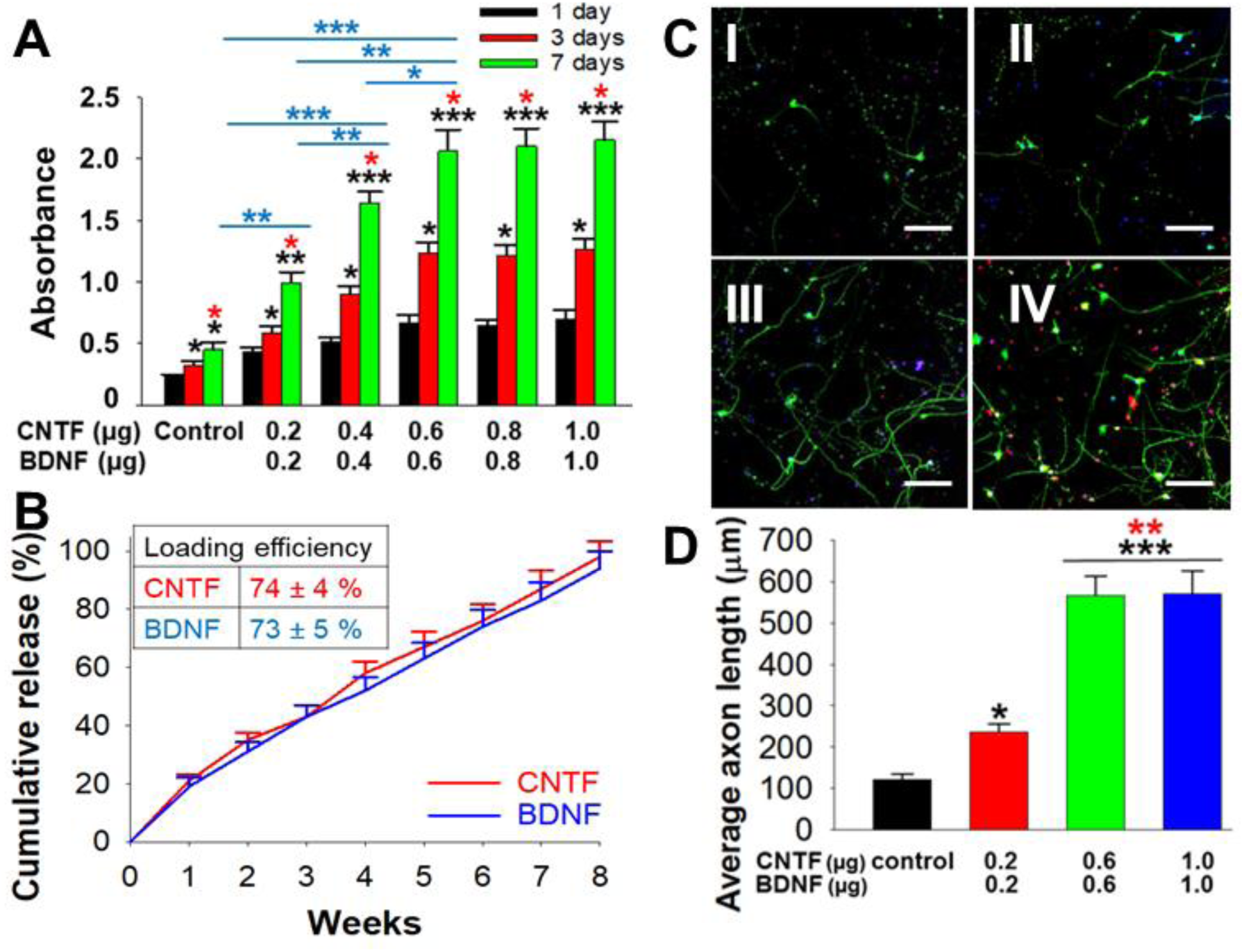
(A) RGC activity depends on different amounts of NTFs. The activity increased significantly up to 0.6 µg of each CNTF and BDNF. However, the amounts over 0.6 µg did not show a further increase in the activity. N=5. *p<0.05, **p<0.01, ***p<0.001 (ANOVA with Bonferroni post-hoc analysis). Error bars indicate standard deviation of the mean. (B) The loading efficiency and the cumulative release of CNTF and BDNF in the absence of DPA. No significant changes were observed compared to the system containing DPA-loaded S-PSHU NPs. N=5. Error bars indicate standard deviation of the mean. (C) The RGC axon growth depends on different amounts of CNTF and BDNF. Tuj-1-stained RGC axons (green) in the group of control (I), 0.2 µg (II), 0.6 µg (III), and 1.0 µg (IV). Scale bar: 200 µm. (D) The bar graph represents the quantification of axon length. Significant axon growth was observed up to 0.6 µg group without further increase in the axon length thereafter. N=5. *p<0.05, **p<0.01, ***p<0.001 (ANOVA with Bonferroni post-hoc analysis). Error bars indicate standard deviation of the mean.

To determine the loading amounts of NTFs, primary RGCs were cultured with NTF- loaded S-PSHU NPs without DPA. The purpose of utilizing DPA is to eliminate excess Zn^2+^ under pathological conditions. The elimination of basal levels of Zn^2+^ from normal (or healthy) RGCs may cause undesired RGC apoptosis. Studies have reported that Zn^2+^ chelation from normal cells resulted in apoptosis across cell types [62–64]. Therefore, DPA was not loaded in the formulation for this study. One concern was whether the absence of DPA in the formulation influences the loading efficiency and release profile of CNTF and BDNF. To address this concern, the same formulation, employed to generate Table 1 and Figure 1, was prepared without DPA for the loading efficiency and release profile. Although slightly increased, the loading efficiency of CNTF and BDNF was very similar to that of NTF-DPA-loaded S-PSHU NPs (Figure 3B, insert). The release profiles of CNTF and BDNF were similar to that observed in Figure 1. Although some variations were observed at each time point, the differences appeared to be minimal. The absence of DPA did not appear to influence the loading efficiency or release profile of CNTF or BDNF.

At 7 days of culture, RGC axon growth was monitored by Tuj-1 staining for the 0.2, 0.6, and 1.0 µg loading groups (Figure 3: C and D). The three groups of NTF-loaded S-PSHU NPs resulted in significantly higher Tuj-1-stained axon growth than the control (p<0.05 for 0.2 µg group and p<0.001 for both 0.6 and 1.0 µg groups). The 0.2 µg group demonstrated an average axon length of 227 µm. The axons in the 0.6 µg group were observed to have an estimated 2.5 times longer axon length than that of the 0.2 µg group (p<0.01). However, the axon length between 0.6 and 1.0 µg groups showed no statistical difference, a similar trend to that of the RGC activity test.

The RGC activity and the axon growth appeared to be positively influenced by increasing amounts of NTFs in the system. There seemed to exist a threshold amount where the effect became capped. In this case, the threshold was observed at 0.6 µg of NTFs.

### S-PSHU NPs preserve the bioactivity of encapsulated NTFs

We observed, in Figure 1 and Figure 3B, that the release of CNTF and BDNF was sustained for 8 weeks. Those release profiles may be beneficial for prolonged RGC survival. Ideally the bioactivity of the encapsulated NTFs should be maintained throughout this period. Previously we reported that a sulfonated delivery matrix has potential for preserving the bioactivity of the encapsulated proteins [15]. Similarly, the capacity of S-PSHU NPs to preserve the bioactivity of the encapsulated NTFs was verified by culturing the primary RGCs with NTF- loaded S-PSHU NPs collected after 4 weeks of release (Figure 4).

**Figure 4.**
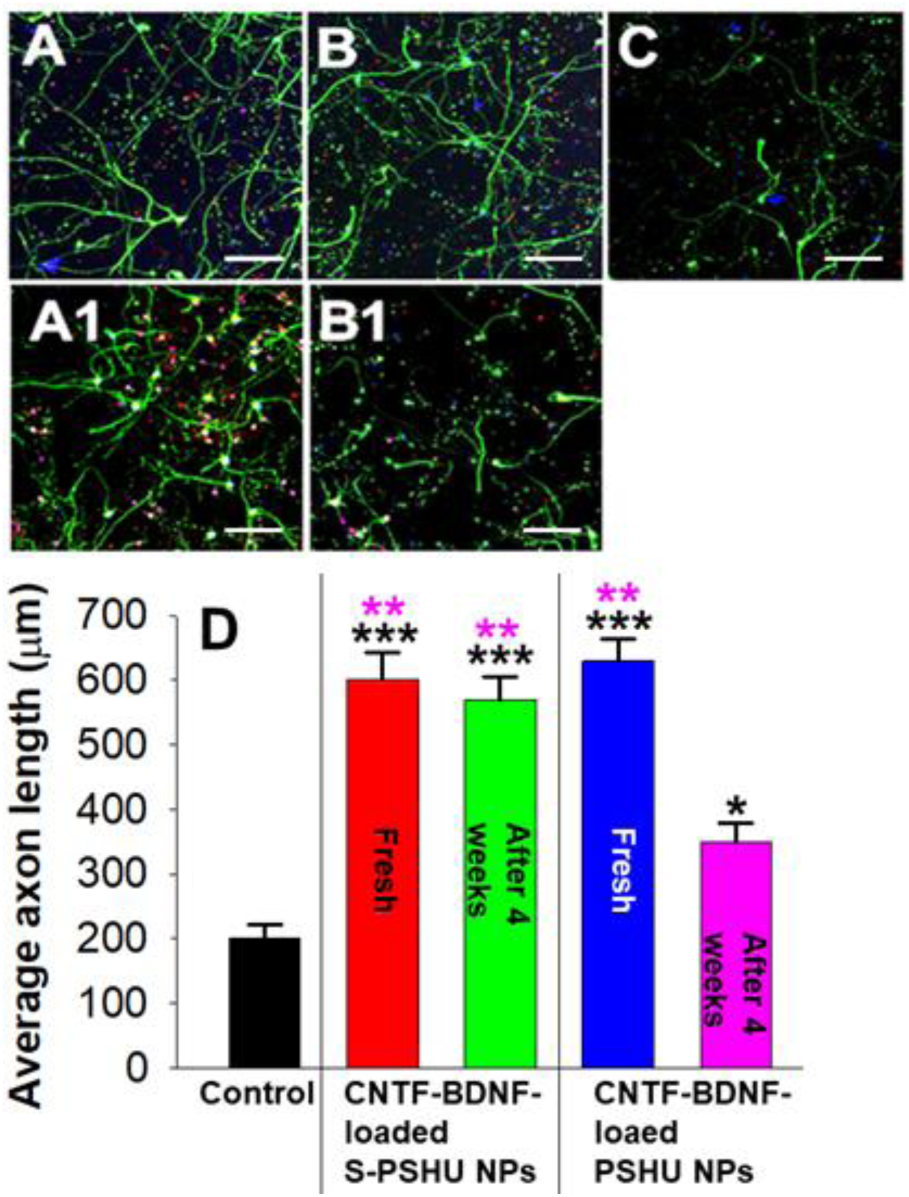
Long-term bioactivity assessment of the encapsulated NTFs. Primary RGCs were treated with NTF-loaded S-PSHU NPs (A and A1), NTF-loaded PSHU NPs (B and B1), and NTF-supported culture medium (C). Scale bar: 200 µm. (D) The bar graph represents the average RGC axon length. The freshly prepared NTF-loaded NPs (A and B) resulted in significantly higher axon growth compared to NTF-supported culture medium. The NTF-loaded S-PSHU NPs collected after 4 weeks of release (A1) resulted a similar axon growth to the freshly prepared NTF-loaded S-PSHU NPs with no statistical difference. The NTF-loaded PSHU NPs collected after 4 weeks of release (B1) showed significantly lower levels compared to the freshly prepared NTF-loaded PSHU NPs and those of S-PSHU NPs. *p<0.05, **p<0.01, ***p<0.001 (ANOVA with Bonferroni post-hoc analysis). N=5. Error bars indicate standard deviation of the mean.

For the groups of S-PSHU NPs (Figure 4: A, A1, and D), the RGC axon growth in the NTF-loaded S-PSHU NPs collected after 4 weeks in PBS showed a slightly decreased trend without statistical difference compared to the freshly prepared NTF-loaded S-PSHU NPs. For the groups of PSHU NPs (Figure 4: B, B1, and D), the freshly prepared NTF-loaded PSHU NPs showed a higher trend in RGC axon growth than the freshly prepared NTF-loaded S-PSHU NPs without statistical difference. However, the axon growth in the NTF-loaded PSHU NPs after 4 weeks in PBS resulted in significantly less RGCs axon growth than the freshly prepared NTF- loaded PSHU NPs (p<0.01), NTF-loaded S-PSHU NPs collected after 4 weeks in PBS (p<0.01), and freshly prepared NTF-loaded S-PSHU NPs (p<0.01). The control group (NTF-supported culture medium, Figure 4C) showed significantly less RGCs axon growth than all other groups. The half-life of native NTFs is known to be short [65, 66]. The bioactivity of NTFs in the control group also seemed to diminish during the culture period, leading to shorter axon extension.

These data proved that the S-PSHU NPs preserved the bioactivity of the encapsulated NTFs for sufficiently long periods of time.

### The NTF-DPA-loaded S-PSHU NPs substantially regenerated RGC axons after ONC

To evaluate treatment efficacy of the system, NTF-DPA-loaded S-PSHU NPs, NTF- loaded S-PSHU NPs or DPA-loaded S-PSHU NPs was prepared using 15 µg of S-PSHU, 0.6 µg of each CNTF and BDNF, and 10 µg of DPA. These loading amounts were selected as no significant benefits were observed thereafter. The retrieved NPs were suspended in 5 µl of saline and intravitreally injected immediately after ONC. At 8 weeks post-intravitreal injection, the CTB-positive (CTB+) axons were monitored to evaluate RGC axon regeneration in the optic nerve sections. The NTF-DPA-loaded S-PSHU NPs resulted in the most extensive CTB+ axons throughout the entire observation field (4.5 mm from the crush site, Figure 5: A, A1, A2, and E), including the number and the length of the CTB+ axons. At each distance from the crush site (marked every 1 mm), the number of CTB+ axons were 2-6 times higher compared to other groups (p<0.001 or p<0.01). The NTF-loaded S-PSHU NPs resulted in higher number of CTB+ axons than DPA-loaded S-PSHU NPs (p<0.01) and saline (p<0.001); however, the number of CTB+ axons were significantly reduced after 3 mm of distance from the crush site, and very few CTB+ axons were observed at 4 mm of distance (Figure 5: B and E). The DPA-loaded S-PSHU NPs resulted in higher number of CTB+ axons than saline (p<0.01); however, it only showed CTB+ axons up to 2 mm of distance from the crush site (Figure 5: C and E). The saline injection showed few CTB+ axons at 1 mm of distance from the crush, which then was negligible thereafter (Figure 5: D and E).

**Figure 5.**
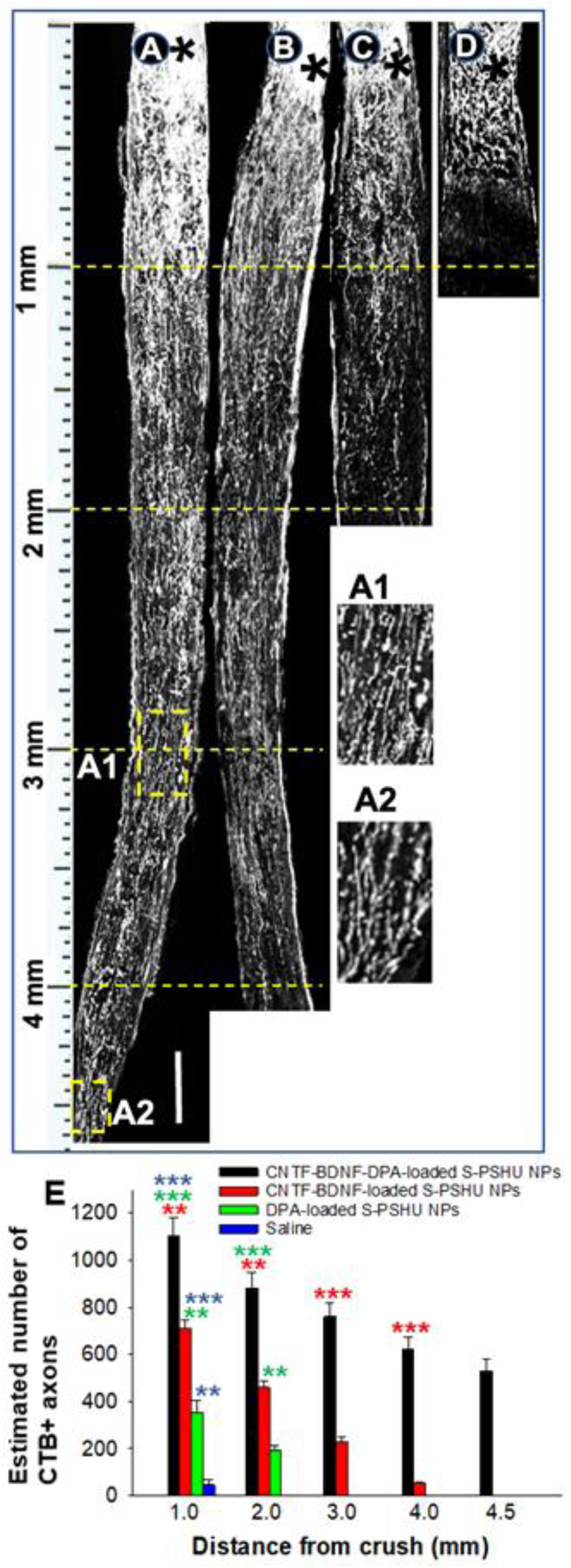
CTB+ axons 8 weeks post-intravitreal injection of NTF-DPA-loaded S-PSHU NPs (A), NTF-loaded S-PSHU NPs (B), DPA-loaded S-PSHU NPs (C), and saline (D) following ONC. A1 and A2 represent magnified images of rectangular regions of A. Scale bar: 350 µm. (E) Estimated number of CTB+ axons at different distances from the crush site. The NTF-DPA-loaded S-PSHU NPs resulted in the most extensive CTB+ axons, which extended throughout the 4.5 mm of entire observation field. The NTF-loaded S-PSHU NPs showed higher number of CTB+ axons than the DPA-loaded S-PSHU NPs. Stars in A, B, C, and D indicate the crush sites. **p<0.01, ***p<0.001 (ANOVA with Bonferroni post-hoc analysis). N=7. Error bars indicate standard deviation of the mean.

### The regenerated axons extended to the distal segment of the brain

In Figure 5, the NTF-DPA-loaded S-PSHU NPs showed the most extensive CTB+ axons throughout the observation field. In addition, the same group expressed CTB+ axons in the proximal optic nerves following an 8-week period (Figure 5, A2), indicating that the CTB+ axons extend beyond the observation field. To investigate whether the CTB+ axons extend to the visual target area in the brain, the expression of CTB+ axons were examined in suprachiasmatic nucleus (SCN, Figure 6: A, B, and C), lateral geniculate nucleus (LGN, Figure 6: D, E, and F), and superior colliculus (SC, Figure 6: G, H, and I) at 8 weeks post-intravitreal injection.

**Figure 6.**
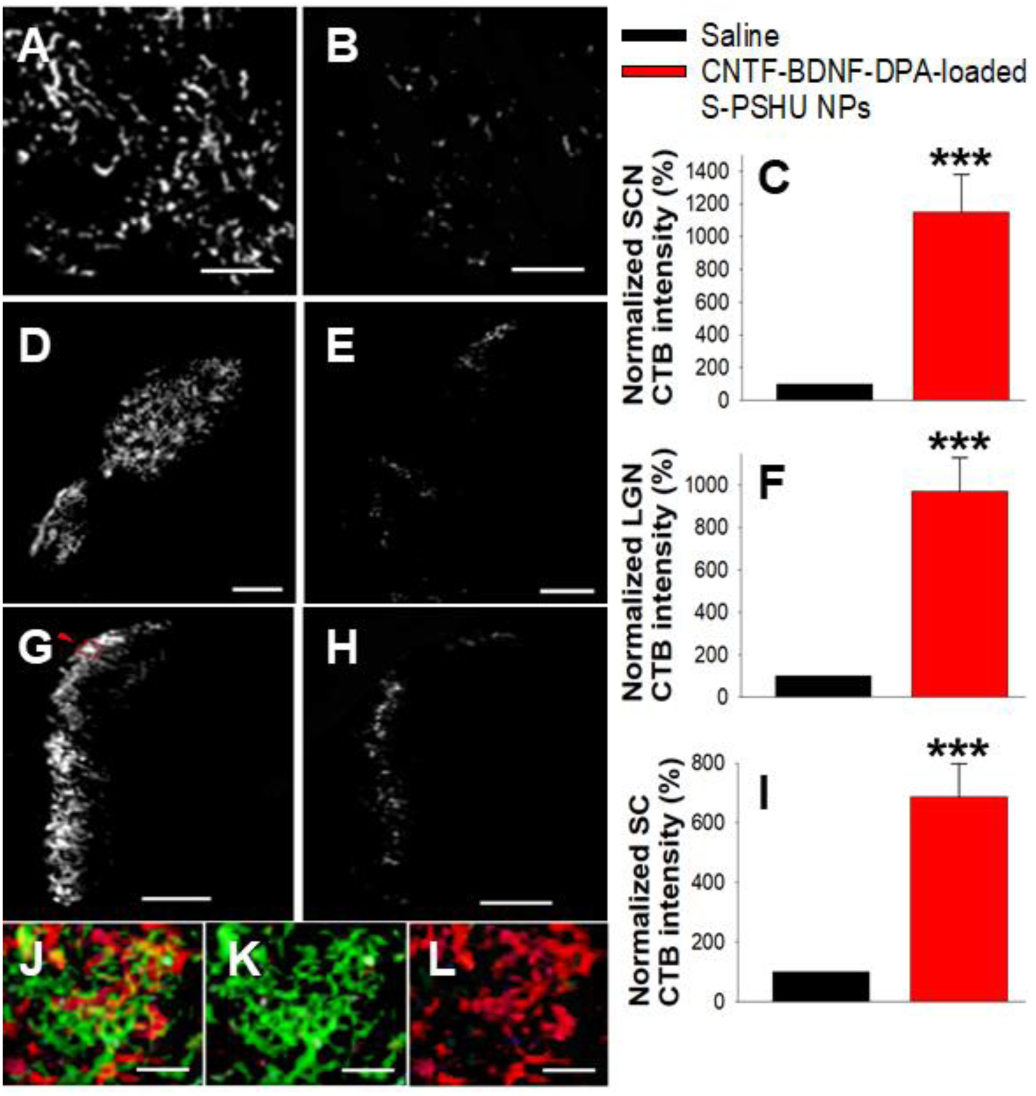
CTB+ axons in SCN (A, B, and C), LGN (D, E, and F), and SC (G, H, and I). At 8 weeks post-intravitreal injection following ONC, NTF-DPA-loaded S-PSHU NPs (A, D, and G) expressed significantly higher CTB+ axons compared to saline (B, E, and H) in all areas. The same group also highly expressed GAP43+ axons in SC (L), indicating that the treatment by NTF-DPA-loaded S-PSHU NPs regenerated axons to visual target areas. Scale bar: 200 µm for A-H; 30 µm for J-L. ***p<0.001 (Student’s t-test). N=20. Error bars indicate standard deviation of the mean.

Although some CTB+ axons were observed in saline group, the expression was very sparse, indicating that most RGC axons degenerated after ONC (Figure 6: B, E, and H). However, the NTF-DPA-loaded S-PSHU NPs resulted in significantly higher CTB+ axons in all areas compared to saline group (Figure 6, A, D, and G, p<0.001). SC is a deeper layer of visual pathway. To demonstrate whether CTB+ axons in this area are nascent or spared, the axons in the red rectangle area of SC (see the red arrow in Figure 6G) were co-labeled with GAP-43 (1:100). The image showed highly expressed GAP43+ axons (Figure 6L), which suggested that the regenerated axons extended to appropriate visual target areas.

### NTF-DPA-loaded S-PSHU NPs significantly improved RGC survival after ONC

At 8 weeks post-intravitreal injection following ONC, RGC survival was evaluated by counting the Tuj-1-stained RGCs on the flat mounted retina and normalizing against those of healthy retina (Figure 7). The NTF-DPA-loaded S-PSHU NPs showed about 83% RGC density compared to those of a healthy retina (Figure 7: A, B and F). Although the RGC survival of this group was lower than the healthy control (p<0.05), it resulted in significantly higher RGC survival than NTF-loaded S-PSHU NPs (64%, p<0.05, Figure 6: C and F), DPA-loaded S-PSHU NPs (42%, p<0.01, Figure 6: D and F), and saline control (10%, p<0.001, Figure 6: E and F). In addition, more structural degeneration of the retina was observed in the lower RGC survival groups. Therefore, co-delivery of NTFs and DPA seemed to support RGC axon regeneration and RGC survival.

**Figure 7.**
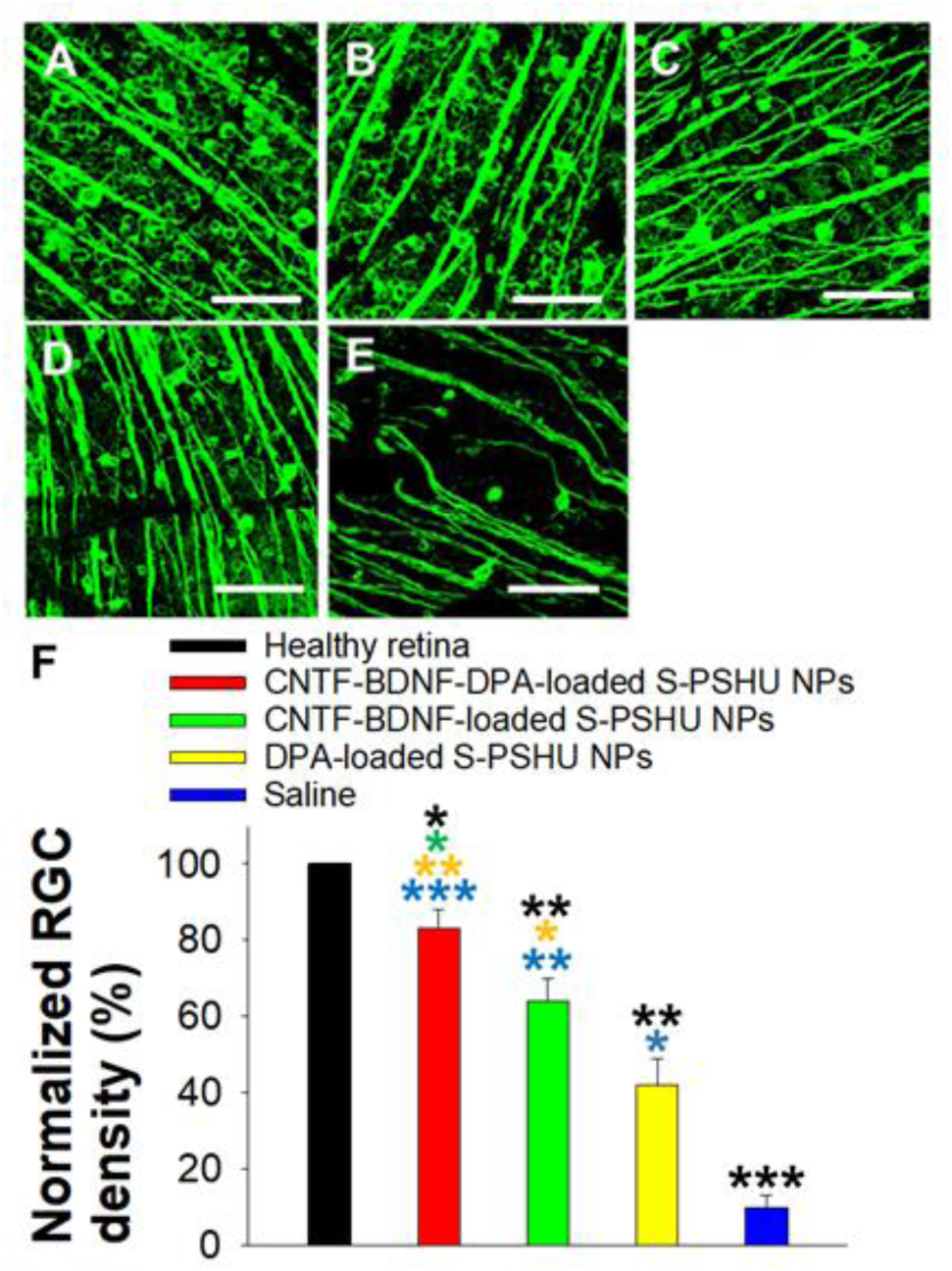
Protective effects of NTF-DPA-loaded S-PSHU NPs against RGC death following ONC. Eight weeks post treatment, retinal flatmounts were immunostained for RGCs using Tuj-1 antibody (green). The representative images of healthy retina (A), NTF-DPA-loaded S-PSHU NPs (B), NTF-loaded S-PSHU NPs (C), DPA-loaded S-PSHU NPs (D), and saline (E) were shown. Scale bar: 100 µm. (F) The RGC density was normalized to that of healthy retina. The NTF-DPA-loaded S-PSHU NPs resulted in the highest RGC survival rate following ONC. The NTF-loaded S-PSHU NPs showed better RGC survival rate than the DPA-loaded S-PSHU NPs. N=10. *p<0.05, **p<0.01, ***p<0.001 (ANOVA with Bonferroni post-hoc analysis). Error bars indicate standard deviation of the mean.

## CONCLUSION

In this study, we aimed to develop a new treatment methodology to improve RGC axon regeneration in traumatic optic neuropathy. To deliver a continuous supply of NTFs and reduce neurotoxicity induced by excessive Zn^2+^, a S-PHU NP-based delivery system was developed to co-deliver NTFs (CNTF and BDNF) and a Zn^2+^ chelator (DPA). *In vitro* release tests demonstrated that the system sustained the release of NTFs and DPA for 8 weeks and 4 weeks, respectively. The sulfonation of NPs enhanced the bioactivity and extended the release of NTFs. This delivery system could eliminate the acutely increased retinal Zn^2+^ after ONC in a DPA-dose dependent manner. When delivering NTFs, an optimal level of loading combination was identified that promotes RGC axon growth *in vitro*. The NTF-DPA-loaded S-PSHU NPs significantly improved RGC survival and axon regeneration in rat retinas following ONC. Regenerated axons were observed to extend into the visual target area in the brain. The NTF- DPA-loaded S-SPHU NPs resulted in significantly higher RGC survival and axon regeneration compared to NTFs alone or DPA alone loaded in S-PSHU NPs, validating the efficacy of this system, which co-addresses the continuous supply of NTFs and the inhibition of Zn^2+^-based neurotoxicity in the traumatic optic neuropathy.

## Acknowledgements

This work was supported by National Institutes of Health (NIH) Grants R01EY031461 (Park); and an Unrestricted Research Grant to the Department of Ophthalmology from Research to Prevent Blindness and CU-Gates Grubstake Award (Nam).

## CRediT authorship contribution statement

Huynh Quang Dieu Nguyen: Data curation, Formal analysis, Writing – review & editing.

Mi-hyun Nam: Formal analysis, Methodology, Writing – review & editing. Jozsef Vigh: Methodology. Joseph Brzezinski: Writing – review & editing. Lucas Duncan: Writing – review & editing. Daewon Park: Conceptualization, Funding acquisition, Methodology, Project administration, Resources, Supervision, Validation, Writing - original draft, Writing - review & editing.

## Data Availability Statement

The data presented in this study are available in the article.

---

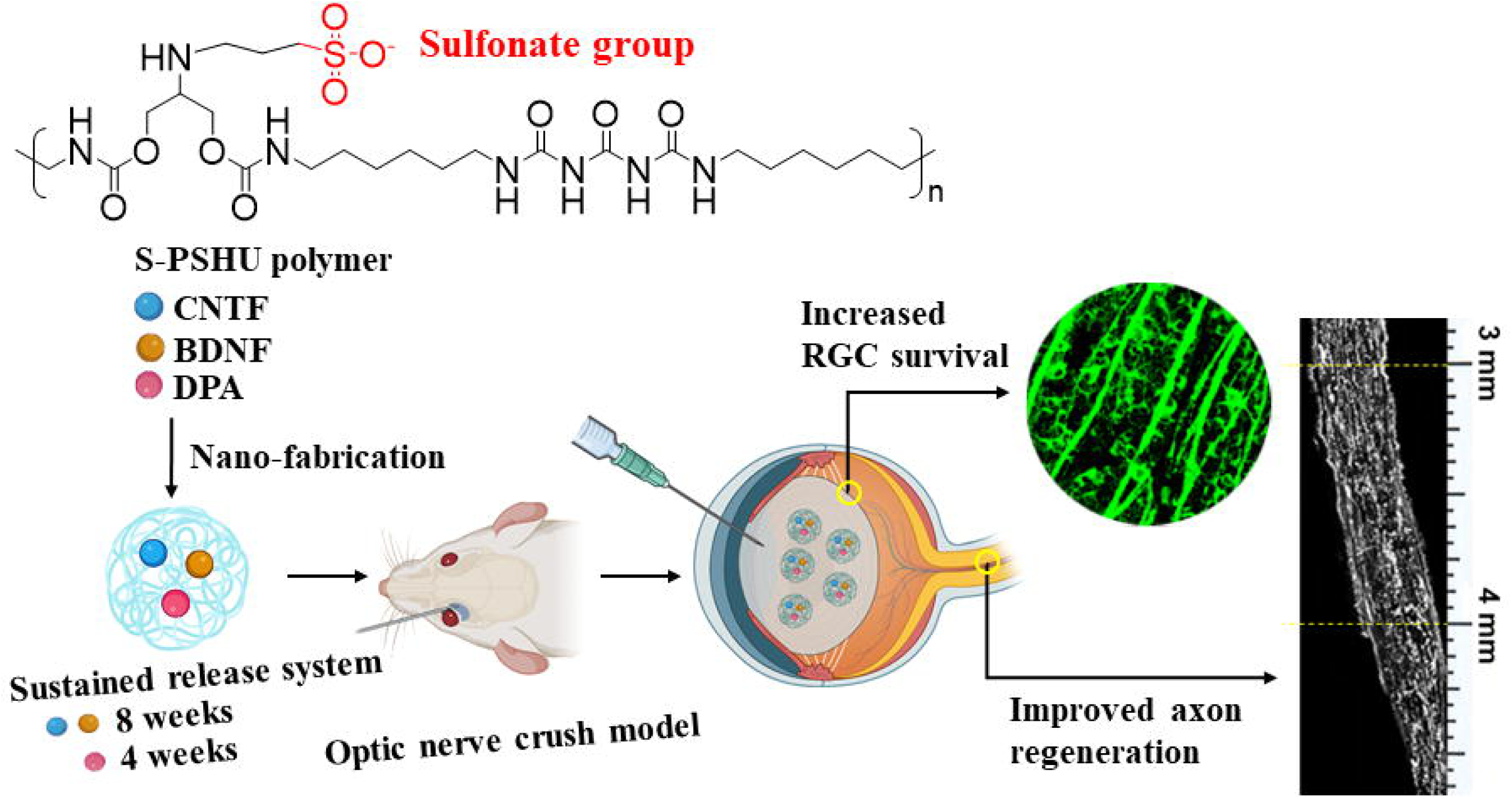

## REFERENCES

[1] N.R. Miller, N.J. Newman, V. Biousse, J.B. Kerrison, Clinical Neuro-Ophthalmology, Second edition ed., Lippincott Williams & Wilkins 2007.

[2] M.L. Ko, D.N. Hu, R. Ritch, S.C. Sharma, The combined effect of brain-derived neurotrophic factor and a free radical scavenger in experimental glaucoma, Invest. Ophthalmol. Vis. Sci. 41(10) (2000) 2967–2971.

[3] G. Parrilla-Reverter, M. Agudo, P. Sobrado-Calvo, M. Salinas-Navarro, M.P. Villegas-Perez, M. Vidal-Sanz, Effects of different neurotrophic factors on the survival of retinal ganglion cells after a complete intraorbital nerve crush injury: A quantitative in vivo study, Exp. Eye Res. 89(1) (2009) 32–41.

[4] M.K. Mathews, Y. Guo, P. Langenberg, S.L. Bernstein, Ciliary neurotrophic factor (CNTF)- mediated ganglion cell survival in a rodent model of non-arteritic anterior ischaemic optic neuropathy (NAION), Br. J. Ophthalmol. 99(1) (2015) 133–137.

[5] M.R. Laughter, J.R. Bardill, D.A. Ammar, B. Pena, D.J. Calkins, D. Park, Injectable Neurotrophic Factor Delivery System Supporting Retinal Ganglion Cell Survival and Regeneration Following Optic Nerve Crush, Acs Biomaterials Science & Engineering 4(9) (2018) 3374–3383.

[6] R.G. Thorne, W.H. Frey, Delivery of neurotrophic factors to the central nervous system - Pharmacokinetic considerations, Clin. Pharmacokinet. 40(12) (2001) 907–946.

[7] B. Mead, A. Logan, M. Berry, W. Leadbeater, B.A. Scheven, Intravitreally Transplanted Dental Pulp Stem Cells Promote Neuroprotection and Axon Regeneration of Retinal Ganglion Cells After Optic Nerve Injury, Invest. Ophthalmol. Vis. Sci. 54(12) 7544–7556.

[8] S.X. Li, B. Hu, D. Tay, K.F. So, H.K.F. Yip, Intravitreal transplants of Schwann cells and fibroblasts promote the survival of axotomized retinal ganglion cells in rats, Brain Res. 1029(1) (2004) 56–64.

[9] Y. Fukuda, M. Watanabe, H. Sawai, T. Miyoshi, Functional recovery of vision in regenerated optic nerve fibers, Vision Res. 38(10) (1998) 1545–1553.

[10] B. Lorber, M. Berry, A. Logan, Different factors promote axonal regeneration of adult rat retinal ganglion cells after lens injury and intravitreal peripheral nerve grafting, J. Neurosci. Res. 86(4) (2008) 894–903.

[11] G. Tezel, X.J. Yang, J.J. Yang, M.B. Wax, Role of tumor necrosis factor receptor-1 in the death of retinal ganglion cells following optic nerve crush injury in mice, Brain Res. 996(2) (2004) 202–212.

[12] P.D. Koeberle, A.K. Ball, Nitric oxide synthase inhibition delays axonal degeneration and promotes the survival of axotomized retinal ganglion cells, Experimental Neurology 158(2) (1999) 366–381.

[13] L. Guo, S.E. Moss, R.A. Alexander, R.R. Ali, F.W. Fitzke, M.F. Cordeiro, Retinal ganglion cell apoptosis in glaucoma is related to intraocular pressure and IOP-induced effects on extracellular matrix, Invest. Ophthalmol. Vis. Sci. 46(1) (2005) 175–182.

[14] Y.Q. Li, L. Andereggen, K. Yuki, K. Omura, Y.Q. Yin, H.Y. Gilbert, B. Erdogan, M.S. Asdourian, C. Shrock, S. De Lima, U.P. Apfel, Y.H. Zhuo, M. Hershfinkele, S.J. Lippard, P.A. Rosenberg, L. Benowitz, Mobile zinc increases rapidly in the retina after optic nerve injury and regulates ganglion cell survival and optic nerve regeneration, Proc. Natl. Acad. Sci. U. S. A. 114(2) (2017) E209–E218.

[15] B. Pena, R. Shandas, D. Park, A heparin-mimicking reverse thermal gel for controlled delivery of positively charged proteins, Journal of Biomedical Materials Research Part A 103(6) (2015) 2102–2108.

[16] M. Almasieh, A.M. Wilson, B. Morquette, J.L.C. Vargas, A. Di Polo, The molecular basis of retinal ganglion cell death in glaucoma, Prog. Retin. Eye Res. 31(2) 152–181.

[17] A.J. Weber, C.D. Harman, BDNF Treatment and Extended Recovery From Optic Nerve Trauma in the Cat, Invest. Ophthalmol. Vis. Sci. 54(10) (2013) 6594–6604.

[18] H. Chen, A.J. Weber, BDNF enhances retinal ganglion cell survival in cats with optic nerve damage, Invest. Ophthalmol. Vis. Sci. 42(5) (2001) 966–974.

[19] D. Fischer, M. Leibinger, Promoting optic nerve regeneration, Prog. Retin. Eye Res. 31(6) (2012) 688–701.

[20] M.H. Chun, W.K. Ju, K.Y. Kim, M.Y. Lee, H.D. Hofmann, M. Kirsch, S.J. Oh, Upregulation of ciliary neurotrophic factor in reactive Muller cells in the rat retina following optic nerve transection, Brain Res. 868(2) (2000) 358–362.

[21] J. Weise, S. Isenmann, N. Klocker, S. Kugler, S. Hirsch, C. Gravel, M. Bahr, Adenovirus-mediated expression of ciliary neurotrophic factor (CNTF) rescues axotomized rat retinal ganglion cells but does not support axonal regeneration in vivo, Neurobiol. Dis. 7(3) (2000) 212–223.

[22] R.E. MacLaren, P.K. Buch, A.J. Smith, K.S. Balaggan, A. MacNeil, J.S. Taylorc, N.N. Osborne, R.R. Ali, CNTF gene transfer protects ganglion cells in rat retinae undergoing focal injury and branch vessel occlusion, Exp. Eye Res. 83(5) (2006) 1118–1127.

[23] Y. Mikata, T. Fujimoto, T. Fujiwara, S.-i. Kondo, Intramolecular ether oxygen coordination in the zinc complexes with dipicolylamine (DPA)-derived ligands, Inorganica Chimica Acta 370(1) (2011) 420–426.

[24] S. Maruyama, K. Kikuchi, T. Hirano, Y. Urano, T. Nagano, A novel, cell-permeable, fluorescent probe for ratiometric imaging of zinc ion, J Am Chem Soc 124(36) (2002) 10650–1.

[25] H.G. Lee, J.H. Lee, S.P. Jang, H.M. Park, S.-J. Kim, Y. Kim, C. Kim, R.G. Harrison, Zinc selective chemosensor based on pyridyl-amide fluorescence, Tetrahedron 67(42) (2011) 8073–8078.

[26] H.G. Lee, J.H. Lee, S.P. Jang, I.H. Hwang, S.-J. Kim, Y. Kim, C. Kim, R.G. Harrison, Zinc selective chemosensors based on the flexible dipicolylamine and quinoline, Inorganica Chimica Acta 394 (2013) 542–551.

[27] Y. Cai, X. Meng, S. Wang, M. Zhu, Z. Pan, Q. Guo, A quinoline based fluorescent probe that can distinguish zinc (II) from cadmium (II) in water, Tetrahedron Letters 54(9) (2013) 1125–1128.

[28] B. Lukowiak, B. Vandewalle, R. Riachy, J. Kerr-Conte, V. Gmyr, S. Belaich, J. Lefebvre, F. Pattou, Identification and purification of functional human beta-cells by a new specific zinc- fluorescent probe, Journal of Histochemistry & Cytochemistry 49(4) (2001) 519–527.

[29] C. Talmard, A. Bouzan, P. Faller, Zinc binding to amyloid-beta: Isothermal titration calorimetry and Zn competition experiments with Zn sensors, Biochemistry 46(47) (2007) 13658–13666.

[30] N. Dinh, A. van der Ent, D.R. Mulligan, A.V. Nguyen, Zinc and lead accumulation characteristics and in vivo distribution of Zn2+ in the hyperaccumulator Noccaea caerulescens elucidated with fluorescent probes and laser confocal microscopy, Environmental and Experimental Botany 147 (2018) 1–12.

[31] J.A.L. Figueroa, K.S. Vignesh, G.S. Deepe, J. Caruso, Selectivity and specificity of small molecule fluorescent dyes/probes used for the detection of Zn2+ and Ca2+ in cells, Metallomics 6(2) (2014) 301–315.

[32] M.R. Karim, D.H. Petering, Newport Green, a fluorescent sensor of weakly bound cellular Zn2+: competition with proteome for Zn2+, Metallomics 8(2) (2016) 201–210.

[33] S.L. Sensi, H.Z. Yin, J.H. Weiss, Glutamate triggers preferential Zn2+ flux through Ca2+ permeable AMPA channels and consequent ROS production, Neuroreport 10(8) (1999) 1723–1727.

[34] R.B. Thompson, D. Peterson, W. Mahoney, M. Cramer, B.P. Maliwal, S.W. Suh, C. Frederickson, C. Fierke, P. Herman, Fluorescent zinc indicators for neurobiology, Journal of Neuroscience Methods 118(1) (2002) 63–75.

[35] M.R. Laughter, D.A. Ammar, J.R. Bardill, B. Pena, M.Y. Kahook, D.J. Lee, D. Park, A Self-Assembling Injectable Biomimetic Microenvironment Encourages Retinal Ganglion Cell Axon Extension in Vitro, Acs Applied Materials & Interfaces 8(32) (2016) 20540–20548.

[36] W.A. Lagreze, A. Pielen, R. Steingart, G. Schlunck, H.D. Hofmann, I. Gozes, M. Kirsch, The peptides ADNF-9 and NAP increase survival and neurite outgrowth of rat retinal ganglion cells in vitro, Invest. Ophthalmol. Vis. Sci. 46(3) (2005) 933–938.

[37] S. Morishita, H. Oku, T. Horie, M. Tonari, T. Kida, A. Okubo, T. Sugiyama, S. Takai, H. Hara, T. Ikeda, Systemic Simvastatin Rescues Retinal Ganglion Cells from Optic Nerve Injury Possibly through Suppression of Astroglial NF-kappa B Activation, Plos One 9(1) (2014).

[38] Q. Cui, H.K. Yip, R.C.H. Zhao, K.F. So, A.R. Harvey, Intraocular elevation of cyclic AMP potentiates ciliary neurotrophic factor-induced regeneration of adult rat retinal ganglion cell axons, Mol. Cell. Neurosci. 22(1) (2003) 49–61.

[39] B.J. Yungher, M. Ribeiro, K.K. Park, Regenerative Responses and Axon Pathfinding of Retinal Ganglion Cells in Chronically Injured Mice, Invest. Ophthalmol. Vis. Sci. 58(3) (2017) 1743–1750.

[40] S. de Lima, Y. Koriyama, T. Kurimoto, J.T. Oliveira, Y.Q. Yin, Y.Q. Li, H.Y. Gilbert, M. Fagiolini, A.M.B. Martinez, L. Benowitz, Full-length axon regeneration in the adult mouse optic nerve and partial recovery of simple visual behaviors, Proc. Natl. Acad. Sci. U. S. A. 109(23) (2012) 9149–9154.

[41] S. Leon, Y.Q. Yin, J. Nguyen, N. Irwin, L.I. Benowitz, Lens injury stimulates axon regeneration in the mature rat optic nerve, J. Neurosci. 20(12) (2000) 4615–4626.

[42] T. Kurimoto, Y.Q. Yin, K. Omura, H.Y. Gilbert, D. Kim, L.P. Cen, L. Moko, S. Kugler, L.I. Benowitz, Long-Distance Axon Regeneration in the Mature Optic Nerve: Contributions of Oncomodulin, cAMP, and pten Gene Deletion, J. Neurosci. 30(46) (2010) 15654–15663.

[43] N. Sangaj, P. Kyriakakis, D. Yang, C.W. Chang, G. Arya, S. Varghese, Heparin Mimicking Polymer Promotes Myogenic Differentiation of Muscle Progenitor Cells, Biomacromolecules 11(12) (2010) 3294–3300.

[44] C.F. Ibanez, Emerging themes in structural biology of neurotrophic factors, Trends in Neurosciences 21(10) (1998) 438–444.

[45] A.M. Adamo, M.P. Zago, G.G. Mackenzie, L. Aimo, C.L. Keen, A. Keenan, P.I. Oteiza, The Role of Zinc in the Modulation of Neuronal Proliferation and Apoptosis, Neurotoxicity Research 17(1) (2010) 1–14.

[46] R.S. Corniola, N.M. Tassabehji, J. Hare, G. Sharma, C.W. Levenson, Zinc deficiency impairs neuronal precursor cell proliferation and induces apoptosis via p53-mediated mechanisms, Brain Research 1237 (2008) 52–61.

[47] H.L. Gao, W. Zheng, N. Xin, Z.H. Chi, Z.Y. Wang, J. Chen, Z.Y. Wang, Zinc Deficiency Reduces Neurogenesis Accompanied by Neuronal Apoptosis Through Caspase-Dependent and - Independent Signaling Pathways, Neurotoxicity Research 16(4) (2009) 416–425.

[48] C.L. Dvergsten, G.J. Fosmire, D.A. Ollerich, H.H. Sandstead, ALTERATIONS IN THE POSTNATAL-DEVELOPMENT OF THE CEREBELLAR CORTEX DUE TO ZINC-DEFICIENCY .2. IMPAIRED MATURATION OF PURKINJE-CELLS, Dev. Brain Res. 16(1) (1984) 11–20.

[49] C.L. Dvergsten, L.A. Johnson, H.H. Sandstead, ALTERATIONS IN THE POSTNATAL- DEVELOPMENT OF THE CEREBELLAR CORTEX DUE TO ZINC-DEFICIENCY .3. IMPAIRED DENDRITIC DIFFERENTIATION OF BASKET AND STELLATE CELLS, Dev. Brain Res. 16(1) (1984) 21–26.

[50] B.K.Y. Bitanihirwe, M.G. Cunningham, Zinc: The Brain’s Dark Horse, Synapse 63(11) (2009) 1029–1049.

[51] C.J. Frederickson, J.Y. Koh, A.I. Bush, The neurobiology of zinc in health and disease, Nature Reviews Neuroscience 6(6) (2005) 449–462.

[52] M.P. Zago, G.G. Mackenzie, A.M. Adamo, C.L. Keen, P.I. Oteiza, Differential modulation of MAP kinases by zinc deficiency in IMR-32 cells: Role of H2O2, Antioxidants & Redox Signaling 7(11-12) (2005) 1773–1782.

[53] A.M. DiGirolamo, M. Ramirez-Zea, Role of zinc in maternal and child mental health, American Journal of Clinical Nutrition 89(3) (2009) 940S–945S.

[54] H. Ra, H.L. Kim, H.W. Lee, Y.H. Kim, Essential role of p53 in TPEN-induced neuronal apoptosis, Febs Letters 583(9) (2009) 1516–1520.

[55] S.W. Suh, S.J. Won, A.M. Hamby, B.H. Yoo, Y. Fan, C.T. Sheline, H. Tamano, A. Takeda, J.L. Liu, Decreased brain zinc availability reduces hippocampal neurogenesis in mice and rats, Journal of Cerebral Blood Flow and Metabolism 29(9) (2009) 1579–1588.

[56] D.W. Choi, J.Y. Koh, Zinc and brain injury, Annual Review of Neuroscience 21 (1998) 347–375.

[57] J.S. Choi, K.A. Kim, Y.J. Yoon, T. Fujikado, C.K. Joo, Inhibition of cyclooxygenase-2 expression by zinc-chelator in retinal ischemia, Vision Research 46(17) (2006) 2721–2727.

[58] M.H. Yoo, J.Y. Lee, S.E. Lee, J.Y. Koh, Y.H. Yoon, Protection by pyruvate of rat retinal cells against zinc toxicity in vitro, and pressure-induced ischemia in vivo, Investigative Ophthalmology & Visual Science 45(5) (2004) 1523–1530.

[59] J.Y. Koh, S.W. Suh, B.J. Gwag, Y.Y. He, C.Y. Hsu, D.W. Choi, The role of zinc in selective neuronal death after transient global cerebral ischemia, Science 272(5264) (1996) 1013–1016.

[60] S.W. Suh, P. Garnier, K. Aoyama, Y.M. Chen, R.A. Swanson, Zinc release contributes to hypoglycemia-induced neuronal death, Neurobiology of Disease 16(3) (2004) 538–545.

[61] S.W. Suh, A.M. Hamby, E.T. Gum, B.S. Shin, S.J. Won, C.T. Sheline, P.H. Chan, R.A. Swanson, Sequential release of nitric oxide, zinc, and superoxide in hypoglycemic neuronal death, Journal of Cerebral Blood Flow and Metabolism 28(10) (2008) 1697–1706.

[62] H.J. Hyun, J.H. Sohn, D.W. Ha, Y.H. Ahn, J.-Y. Koh, Y.H. Yoon, Depletion of Intracellular Zinc and Copper with TPEN Results in Apoptosis of Cultured Human Retinal Pigment Epithelial Cells, Invest. Ophthalmol. Vis. Sci. 42(2) (2001) 460–465.

[63] B. Zhu, J. Wang, F. Zhou, Y. Liu, Y. Lai, J. Wang, X. Chen, D. Chen, L. Luo, Z.-C. Hua, Zinc Depletion by TPEN Induces Apoptosis in Human Acute Promyelocytic NB4 Cells, Cellular Physiology and Biochemistry 42(5) (2017) 1822–1836.

[64] M.J. McCabe, S.A. Jiang, S. Orrenius, CHELATION OF INTRACELLULAR ZINC TRIGGERS APOPTOSIS IN MATURE THYMOCYTES, Laboratory Investigation 69(1) (1993) 101–110.

[65] F. Dittrich, H. Thoenen, M. Sendtner, CILIARY NEUROTROPHIC FACTOR - PHARMACOKINETICS AND ACUTE-PHASE RESPONSE IN RAT, Annals of Neurology 35(2) (1994) 151–163.

[66] J.Y. Cai, F.Z. Hua, L.H. Yuan, W. Tang, J. Lu, S.C. Yu, X.F. Wang, Y.H. Hu, Potential Therapeutic Effects of Neurotrophins for Acute and Chronic Neurological Diseases, Biomed Research International (2014).

[67] J.H. Lee, S. Lee, Y.S. Gho, I.S. Song, H. Tchah, M.J. Kim, K.H. Kim, Comparison of confocal microscopy and two-photon microscopy in mouse cornea in vivo, Exp. Eye Res. 132 (2015) 101–108.

