## Supplement document for "Co-delivery of neurotrophic factors and a zinc chelator substantially promotes axon regeneration in the optic nerve crush model"

**Isolate retinal ganglion cells (RGCs)**

RGCs were isolated from rat pups (postnatal day 5−7). The enucleated eyes were transferred to a Petri dish filled with D-PBS. A small incision was made along the anterior part of the eye. Using tweezers, the eye was pulled along this incision line. At this point, the retina was peeled away from the sclera. The dissected retinas were dissociated using the Neural Tissue Dissociation Kit (Miltenyi Biotec). Then, the RGCs were purified using the Retinal Ganglion Cell Isolation Kit (Miltenyi Biotec).

**Determine RGCs axon length**

Z-stack projections were sampled using a 20x objective. The simple neurite tracer plugin (FIJI) was used to analyze the axon length. Each neurite was traced from the cell body to the end of the image frame. The average neurite length was determined by measuring the total length of all axons in three dimensions and dividing this number by the total cell count within that field.

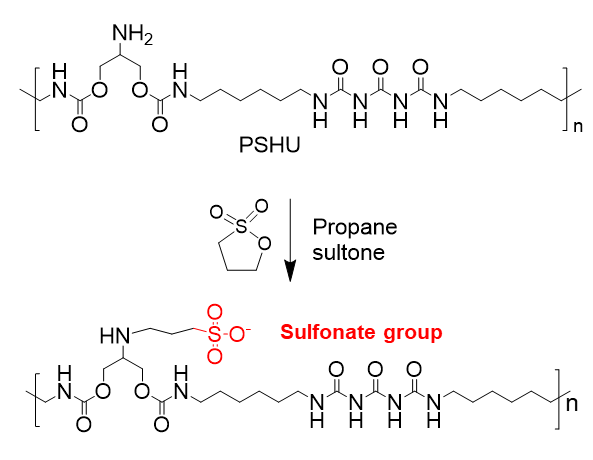

Figure S1. Synthesis of S-PSHU. The amine groups in PSHU were sulfonated using propane sultone to synthesize S-PSHU.

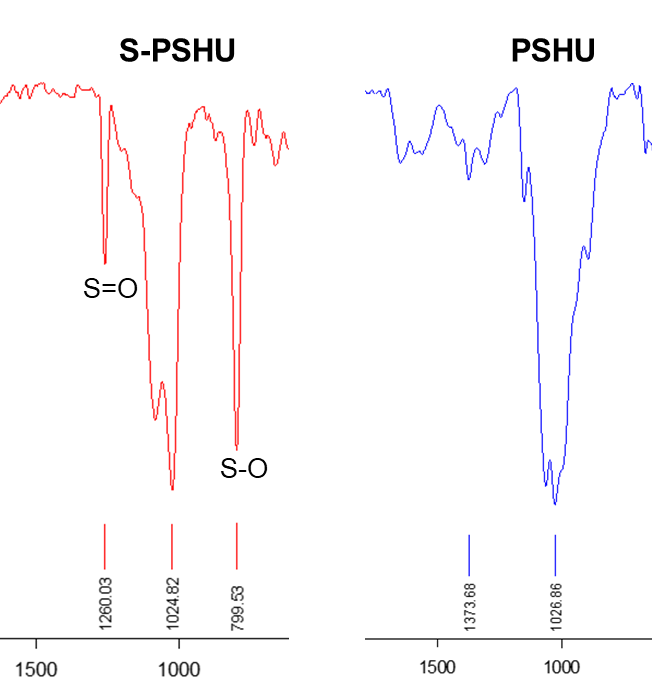

Figure S2. FTIR spectra of S-PSHU and PSHU. After sulfonation of PSHU, S=O and S-O stretching were observed at 1,260 cm^-1^ and 799 cm^-1^, respectively, in S-PSHU.

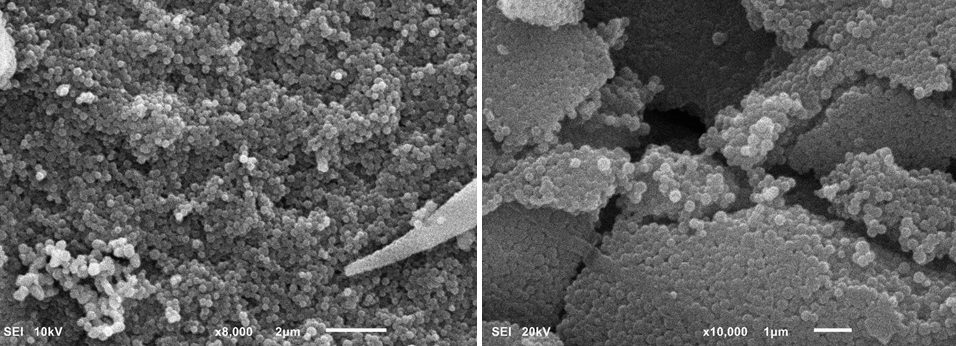

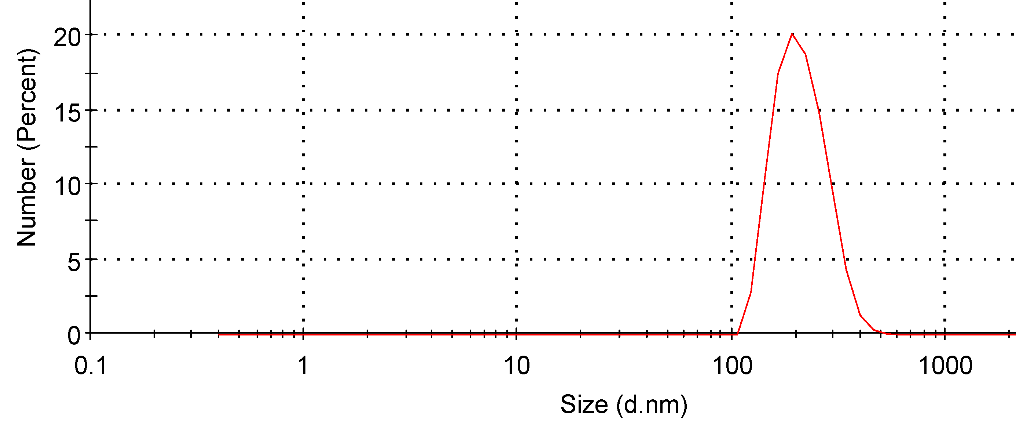

Figure S3. SEM images of CNTF-BDNF-DPA-loaded S-PSHU-NPs (**Top**) and size distribution of NPs by zeta sizer (**Bottom**). The images showed uniform size of NPs with 223±16 nm of diameter.
